# Shuttling of Dorsal by Cactus: mechanism and implications

**DOI:** 10.1101/739284

**Authors:** Allison E. Schloop, Sophia Carrell-Noel, Gregory T. Reeves

**Affiliations:** Genetics Program, North Carolina State University, Raleigh, NC; Department of Chemical and Biomolecular Engineering, North Carolina State University, Raleigh, NC

**Keywords:** Dorsal, Cactus, Toll, shuttling, diffusion, embryo, morphogen, dorsal-ventral axis, development

## Abstract

In a developing animal, morphogen gradients act to pattern tissues into distinct domains of cell types. However, despite their prevalence in development, morphogen gradient formation is a matter of debate. In our recent publication, we showed that the Dorsal/NF-κB morphogen gradient, which patterns the DV axis of the early Drosophila embryo, is partially established by a mechanism of facilitated diffusion. This mechanism, also known as “shuttling,” occurs when a binding partner of the morphogen facilitates the diffusion of the morphogen, allowing it to accumulate at a given site. In this case, the inhibitor Cactus/IκB facilitates the diffusion of Dorsal/NF-κB. In the fly embryo, we used computation and experiment to not only show that shuttling occurs in the embryo, but also that it enables the viability of embryos that inherit only one copy of *dorsal* maternally. Here we further discuss our evidence behind the shuttling mechanism, the previous literature data explained by the mechanism, and how it may also be critical for robustness of development. Finally, we describe an interaction between Dorsal and BMP signaling that is likely affected by shuttling.

## Introduction

Animal development is orchestrated by morphogen gradients which help to pattern tissues based on location and concentration. The formation of these gradients has been a subject of intense scrutiny [1–3]. In *Drosophila*, Dorsal (Dl), a maternally provided transcription factor, acts as a morphogen to regulate patterning of the dorsal-ventral (DV) axis of the early (1-3 hr old) embryo [4]. During these stages of embryogenesis, the nuclei rapidly divide without cytokinesis. These nuclei share a common cytoplasm, and thus, intracellular proteins may diffuse from one nucleus to another. The first nine nuclear division cycles after fertilization occur within the volume of the embryo. After nuclear cycle (nc) 9, the nuclei migrate to the periphery of the embryo, forming a syncytial blastoderm. Four further nuclear divisions occur, and during nc 14, cell membranes introgress between the nuclei in a process called cellularization. Roughly 45 minutes after the start of nc 14, gastrulation takes place, after which, Dl signaling ceases.

While initially uniformly distributed, a nuclear concentration gradient of Dl begins to develop during nuclear cycle (nc) 10. This ultimately results in high concentrations in ventral nuclei and a spatial gradient that decreases to basal concentrations in dorsal nuclei [4–9]. Establishment of this gradient leads to the activation of different genes around the DV axis depending on the concentration of Dl [10,11]. While in the cytoplasm, Dl is bound to Cactus (Cact), which prevents movement into the nucleus; degradation of Cact through Toll signaling on the ventral side releases Dl from Cact and allows it to enter [10,12–15]. Taken together, this information suggests that a depletion of Dl in the cytoplasm around the ventral-most nuclei would occur, creating a cytoplasmic Dl gradient opposite to that of nuclear Dl [4,16]. However, recent work indicates that there is an accumulation of total Dl (nuclear Dl+cytoplasmic Dl) on the ventral side of the embryo as development progresses (Fig. 1) [5,17]. The current paradigm -- Toll signaling, the degradation of Cact, and the local nuclear import of Dl -- cannot explain this accumulation. Therefore, through both computational and experimental means, we have suggested that a facilitated diffusion, or shuttling, mechanism is responsible for the accumulation of total Dl with Cact acting as a carrier molecule [17]. In such a mechanism, a carrier molecule, in this case Cact, facilitates the diffusion of the primary molecule (Dl) against its own concentration gradient.

**Figure 1:**
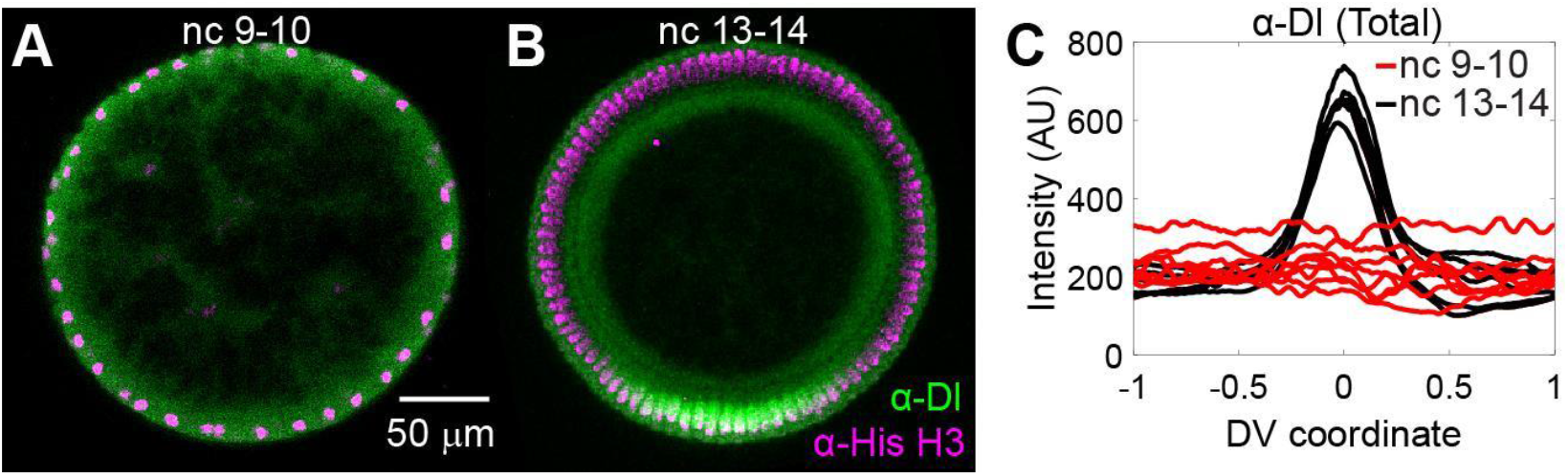
Accumulation of Dl on the ventral side. (A) In nuclear cycle (nc) 9-10 embryos, Dl protein (green) is distributed uniformly around the embryo. (B) In older embryos (nc 13-14), Dl protein has accumulated on the ventral side. (C) Quantification of the total Dl fluorescence intensity (nuclear + cytoplasmic) in nc 91-0 vs nc 13-14 embryos. Magenta in A and B: Histone H3 protein to mark the nucleus. Modified with permission from [17].

Shuttling mechanisms have already been described in conjunction with the formation of signaling gradients in the *Drosophila* embryo; two examples include gradient formation in the Dpp/BMP signaling pathway and establishment of the Spätzle (Spz) gradient. In the Dpp/BMP signaling pathway, the shuttling mechanism, in which an inhibitory complex of Short gastrulation (Sog) and Twisted gastrulation (Tsg) facilitates transport of the BMP ligand Dpp to the dorsal region of the embryo, is well established [18–25]. Dpp signaling begins early in nc 14 as a weak, broad distribution on the dorsal half of the embryo, but becomes highly intense and concentrated at the dorsal midline by late nc 14 [24]. Laterally expressed Sog and Tsg form a complex with Dpp and a second BMP ligand, Scw. The complex prevents Dpp signalling, prohibits movement away from the dorsal region, and protects Dpp from degradation. Loss of either Sog or Tsg results in a failure to accumulate Dpp signaling along the dorsal midline [20]. Tolloid, a metalloprotease that cleaves Sog to release active BMP ligand, is expressed on the dorsal half of the embryo, and is required for shuttling [21].

For Spz, the Toll ligand, a theoretical self-organized shuttling mechanism has been proposed [16] in which Spz is cleaved into two parts that rejoin to form a complex. This complex can experience several outcomes, one of which is dissociation into an inhibitory molecule (N-Spz) and an active molecule (C-Spz). Through modeling work, Haskel-Ittah et al. have suggested that these two molecules can reassociate to form an inactive complex. This complex freely diffuses around the DV axis, but upon reaching the ventral region of the embryo, the N-Spz of this inactive complex is degraded. This mechanism would cause an accumulation of the active molecule, C-Spz, on the ventral side.

The shuttling mechanism is not limited to *Drosophila*. In *Xenopus* and *Danio*, the Sog homolog Chordin performs a similar role in concentrating BMP activity to the ventral side of these embryos [26]. This evidence indicates that not only is shuttling important for the establishment of multiple morphogen gradients, it also occurs during gradient establishment in many organisms.

## Methods

### Fly lines

For maternal over expression of the Mad and Medea constructs, we used a matα-Gal4 line available from the Drosophila Bloomington Stock Center (w[*]; P{w[+mC] = matalpha4-GAL4-VP16}V37, BS#7063). We are grateful to Laurel Raftery for kindly providing the UAS-Med and UAS-Mad lines [27].

### Fluorescent antibody staining

Standard protocols were followed for fixation of 2-4 hr embryos in 37% formaldehyde solution [28].Fluorescent antibody staining was performed according to standard protocols [28], with primary antibodies against Dl (1:10 dilution; DSHB anti-dorsal 7A4) and histone (1:5000 dilution; abcam ab1791). The anti-Dl monoclonal antibody (DSHB Hybridoma Product anti-dorsal 7A4) was deposited to the DSHB by R. Steward [15], and was obtained from the Developmental Studies Hybridoma Bank, created by the NICHD of the NIH and maintained at The University of Iowa, Department of Biology, Iowa City, IA 52242. Secondary antibodies used were donkey anti-mouse-488 (1:500 dilution; Invitrogen A21202) and donkey anti-rabbit-546 (1:500 dilution; Invitrogen A10040),

### Mounting and imaging embryos

Previously-published protocols for mounting and imaging embryos were used [29]. Briefly, fixed embryos were manually cross sectioned in 70% glycerol using a razor blade. The anterior and posterior 1/3rds of the embryo were removed, and the remaining 1/3 (trunk portion of the embryo) was mounted vertically on a microscope slide. Two pieces of double-sided tape were used on either side of the embryos to create a vertical space so that the coverslip would not deform the embryos. Embryos were imaged using a Zeiss 710 laser scanning confocal microscope. 10-20 z-slices were taken for each embryo.

### Image analysis

Image analysis was performed according to our previously published protocols [30]. Briefly, the nuclei were segmented, and the Dl intensity in the nuclei was measured. Gradient width was determined by fitting the Dl gradient to a Gaussian-like shape [5,6,30]. For figures 2C and 3A,B, each z-slice was normalized independently to accurately depict the shape of the gradient.

### Evidence of shuttling in the Dorsal gradient

The Dl/Cact system follows the four features necessary for a shuttling mechanism to occur (see Fig. 2A) [17,31]: (1) Dl, the primary molecule, binds to Cact, a carrier molecule, preventing its capture or degradation; (2) the complex of Dl/Cact can diffuse on a global scale; (3) the complex is degraded in a spatially-dependent manner, which for Dl/Cact is through Toll signalling on the ventral side; and (4) once free of its carrier, the primary molecule can be captured (in this case, by the nucleus) or degraded. The fourth feature prevents free cytoplasmic Dl from diffusing back to the dorsal side (counter-diffusion). Any system with these four features, balanced in the proper way, may exhibit shuttling. Indeed, previous models of the Dl/Cact system, which captured these four features, exhibited shuttling without the express intention of the modelers [32].

**Figure 2:**
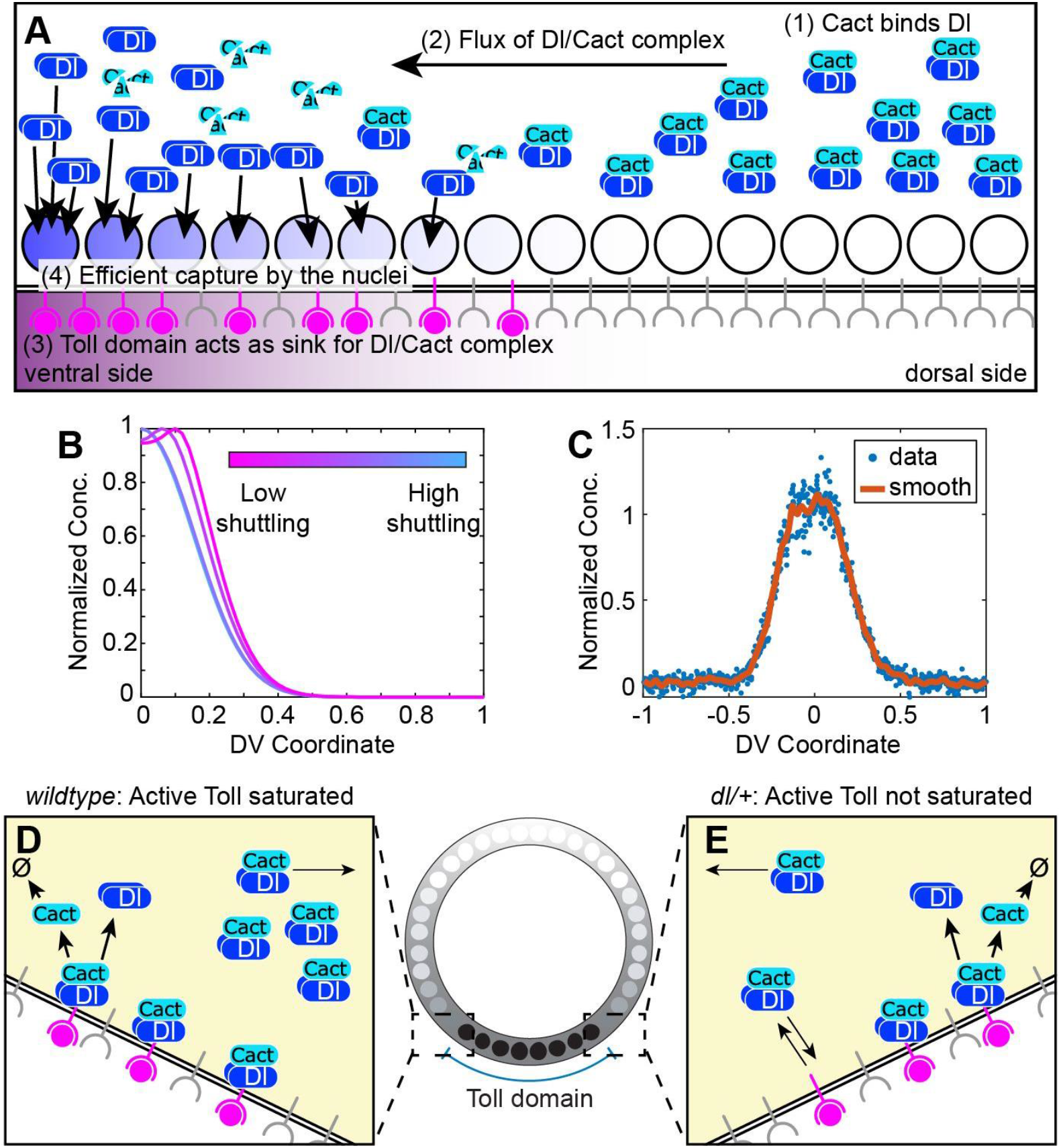
Description of shuttling mechanism..(A) Four biophysical processes necessary for shuttling to occur. Note that process (4), capture by the nuclei, is necessary to prevent free Dl in the cytoplasm from diffusing back to the dorsal side. (B) An example of the hallmark phenotypes that occur when the strength of shuttling is lowered: the gradient becomes wider, flatter on top, or splits into two peaks. (C) Typical embryo from mother heterozygous null for *dl*. The Dl gradient is flatter on top, and is also wider. (D) In wildtype embryos, Toll saturation likely occurs in the tail of the Toll domain. Sufficient numbers of Dl/Cact complexes saturate the active Toll signaling complex, which allows other Dl/Cact complexes to shuttle past towards the ventral midline. (E) In *dl*/+ embryos, there are fewer Dl/Cact complexes, so that active Toll is less saturated, and a lower fraction of Dl/Cact complexes shuttle to the ventral midline. Modified with permission from [17].

Each of these features is well established in the Dl/Cact system -- including point (2), which we confirmed by live imaging of photoactivatable GFP-tagged Dl; [17] -- yet it remained to be seen whether these four processes were balanced in such a way that shuttling occurred in an appreciable manner. Our model of the Dl/Cact system suggested that, if shuttling did occur appreciably, then decreasing the ventrally-directed flux due to shuttling would result in one of a hallmark set of phenotypes, depending on the degree to which the flux lowered [17]. By slowing the diffusion of Dl and Dl/Cact complex, the Dl gradient would widen, become flat on top, or (most severely) split into two peaks (Fig. 2B). In a typical morphogen gradient system (without shuttling), in which the morphogen is produced locally and spreads, lowering the diffusion is expected to narrow the gradient. In contrast, in a shuttling system, diffusion occurs against the gradient of the morphogen, and thus, lowering diffusion would widen the gradient and prevent proper peak accumulation. In a series of experiments in which Dl was tagged with bulky molecules, we found a correlation between widening of the Dl gradient and the degree to which the bulky tags were expected to lower Dl/Cact diffusion [17]. While widening of the gradient is the weakest of the predicted hallmark phenotypes of shuttling, the correlation was telling.

In addition to reduced diffusivity resulting in a hallmark shuttling phenotypes, it was also shown that embryos from heterozygous *dl* null mothers exhibited the hallmark phenotypes to varying degrees (Fig. 2C) [6,17,32]. However, no previous model of the Dl gradient, including our initial models, could reproduce this phenotype when lowering only the dosage of Dl [17,32–34]. The Dl gradients from each of the models were predicted to globally scale with dosage (a half dose of maternal *dl* resulted in a 50% reduction of the gradient, everywhere), so that changes in shape, including widening or flattening, were never observed. On the other hand, we discovered that if the concentration of active Toll receptors were sufficiently low such that they could be saturated by Dl/Cact complexes, the change in gradient shape could possibly be explained [17]. In wildtype embryos, saturation in lateral regions allows accumulation of Dl at the ventral midline and normal gradient formation (Fig. 2D). However, when maternal *dl* production is reduced *(dl* heterozygous embryos), unsaturated Toll receptors trap what limited Dl/Cact complexes are available, preventing accumulation and normal gradient formation (Fig. 2E). The altered Dl gradient shape in *dl* heterozygous embryos is partially restored to the Gaussian-like shape when *Tl* levels are reduced [17].

Hallmark shuttling phenotypes have also been previously observed in several other, seemingly unrelated instances. These include when the domain of active Toll receptors is widened and when the Dl gradient is ectopically expressed along the anterior-posterior axis [17]. In each of these cases, the combination of shuttling and Toll saturation is adequate to explain these observations.

Beyond the hallmark phenotypes observed when the Dl gradient is measured, the shuttling mechanism also allows for intuition regarding a litany of literature observations that were previously unexplained. For example, *dl* transgene fusions to *lacZ* are antimorphic, which is readily explained by the Dl-βgalactosidase forming bukly groups by self-association [17,35]. As another example, a greatly expanded domain of active Toll results in the duplication of the ventral furrow at gastrulation [36]; this arises as an extreme instance of the split peak phenotype [17].

### Importance of shuttling

The evidence for shuttling and Toll saturation strongly suggest the four features necessary for shuttling are indeed balanced such that Dl shuttling occurs *appreciably* in the embryo. On the other hand, given that asymmetric Toll signaling alone, without shuttling, is sufficient to explain the development of a strong ventral-to-dorsal bias in the Dl nuclear concentration, it raises the question of whether a shuttling mechanism is *required* for proper gene expression in the wildtype embryo. If so, then it stands to reason that *cact* function would be necessary to accumulate Dl to the peak levels required for the expression of the ventral-most genes, such as *sna* and *twi*. However, seminal work studying Cact function found that the *twi* domain is expanded in embryos from *cact* null mothers [37]. On the other hand, recent quantitative measurements of the Dl gradient in embryos from mothers with severe loss of *cact* function are consistent with shuttling being required for proper *sna* expression [38]. Further quantitative study of the precise distribution of Dl in *cact* mutants is required to answer this question.

Another question that may arise is whether shuttling is important in other cases, such as to make the Dorsal gradient more robust. Our findings suggest that shuttling is indeed required to accumulate peak levels of Dl in embryos from mothers heterozygous for *dl*. In embryos with compromised shuttling, in the background of maternally heterozygous *dl*, the Dl gradient is severely perturbed in shape and width. Furthermore, the *sna* expression domain is either absent or narrowed to the point of loss of embryonic viability [17].

In addition, one of the hallmark phenotypes of shuttling, in which the Dl gradient splits into two peaks, has been found in the context of other perturbations. This may imply that shuttling provides a role in those processes. For example, when investigating the scaling properties of the Dl gradient, a split-peak of the Dl nuclear gradient was found in the largest embryos (Fig. 3A) [39]. Also, *sna* expression boundaries in these embryos did not scale with size. One possibility is that shuttling allows for size-dependent scaling of the Dl gradient within certain ranges; if the embryo size exceeds certain limits, the shuttling-dependent scaling mechanism fails.

**Figure 3:**
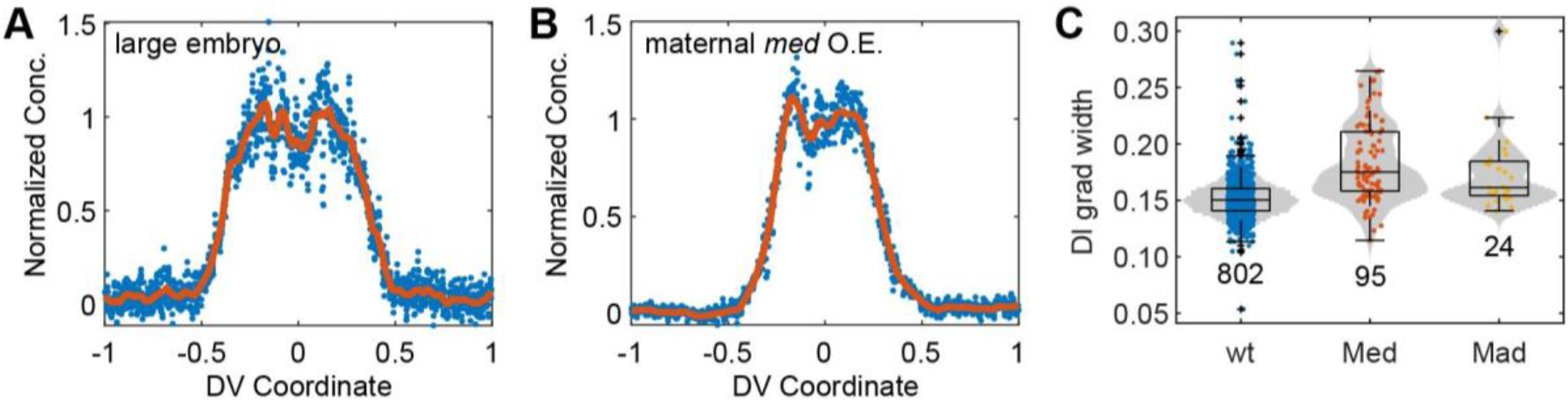
Shuttling phenotypes in other contexts. (A) Quantification of the Dl gradient in embryos with very large DV axes reveals a split-peaked embryo. It is possible that shuttling is responsible for size-dependent scaling of the Dl gradient, which breaks down when the DV axis becomes too large. Modified from [39]. (B) The hallmark phenotypes of shuttling also appear in embryos that have maternally overexpressed components of the BMP pathway. (C) Box-and-violin plot of measure of the Dl gradient width in wildtype, Med-GFP O.E., and Mad-GFP O.E. embryos. Both the Med- and Mad-GFP O.E. embryos have statistically wider gradients (p-val < 10^-150^).

Furthermore, we have found that maternally over-expressing components of the Dpp pathway results in hallmark shuttling phenotypes. We used a maternal Gal4 driver to overexpress the BMP signal transduction components Mad or Medea (see Methods); the resulting embryos had widened Dl gradients, while some had flat tops or double peaks (Fig. 3B,C). Thus, it is possible that the shuttling mechanism also somehow mediates the interactions of the maternal Dpp pathway with the Dl gradient [38,40–43].

## Conclusions

The shuttling phenomenon carries with it many counter-intuitive observations, such as a protein classically construed as an inhibitor that can also potentiate the highest levels of signaling [19–21,23,24,38]. Even so, the phenomenon of shuttling arises naturally when only a handful of sufficient conditions are met: an inhibitor must bind, the complex must be able to diffuse, there must be a sink for the complex, and the shuttled molecule must not participate in significant counter-diffusion. Each of these processes are known to operate in the early *Drosophila* embryo for the Dl/Cact system, and are balanced in such a way that shuttling of Dl by Cact occurs appreciably.

The shuttling mechanism for the Dl gradient appears to be a modest revision of our understanding of Dl gradient formation; however, the mechanism is impactful in many ways. It explains how the Dl gradient levels in the ventral-most nuclei continually grow over time, as well as a litany of other previously unexplained literature observations. Furthermore, it appears to be a crucial mechanism for the robustness of embryo development with respect to several common perturbations, including maternal *dl* dosage compensation, size-dependent scaling, and interactions with other signaling pathways. Future work on the Dl gradient may uncover still more perturbations with phenotypes consistent with the hallmarks of shuttling.

## Acknowledgements

We thank Dr. Thomas Jacobsen for initial discussions of the manuscript.

